# Direct brain recordings reveal continuous encoding of structure in random stimuli

**DOI:** 10.1101/2021.10.01.462295

**Authors:** Julian Fuhrer, Kyrre Glette, Jugoslav Ivanovic, Pål Gunnar Larsson, Tristan Bekinschtein, Silvia Kochen, Robert T. Knight, Jim Tørresen, Anne-Kristin Solbakk, Tor Endestad, Alejandro Blenkmann

## Abstract

The brain excels at processing sensory input, even in rich or chaotic environments. Mounting evidence attributes this to the creation of sophisticated internal models of the environment that draw on statistical structures in the unfolding sensory input. Understanding how and where this modeling takes place is a core question in statistical learning and predictive processing. In this context, we address the role of transitional probabilities as an implicit structure supporting the encoding of a random auditory stream. Leveraging information-theoretical principles and the high spatiotemporal resolution of intracranial electroencephalography, we analyzed the trial-by-trial high-frequency activity representation of transitional probabilities. This unique approach enabled us to demonstrate how the brain continuously encodes structure in random stimuli and revealed the involvement of a network outside of the auditory system, including hippocampal, frontal, and temporal regions. Linking the frameworks of statistical learning and predictive processing, our work illuminates an implicit process that can be crucial for the swift detection of patterns and unexpected events in the environment.

---

Efficient encoding of patterns in ongoing sensory input is critical for survival in an ever-changing environment. Pattern encoding involves the continuous updating of internal representations of the environment based on statistical structures derived from the sensory signal (1–7). The brain is not inherently aware of the underlying structures in the environment and potential regularities in the sensory stream must be assessed with regard to previously encoded regularity (8–10). Sensitivity to conditional regularity between events has been observed in humans (11–21) and animals (22, 23). Because events in the environment rarely occur independently, this pattern extraction is necessary for the fast and efficient processing of sensory information.

A mathematical representation of such conditional regularity is transitional probabilities (TPs). TPs describe how likely one event predicts another. That is the ratio of the directional co-occurrence of events given their frequency (3, 24–26). As an example, experimental studies in infants and adults have shown that the TPs between syllables constitute patterns that facilitate the identification of word-like units (11, 26–30), thus making TP encoding essential for language development (3, 4, 25, 28, 31–33).

While the brain’s sensitivity to conditional regularities has been observed in experimental studies across sensory domains, the underlying mechanisms remain poorly understood (3, 27, 28, 34–41). Studies on sensory processing and statistical learning have reported engagement of multiple brain structures, suggesting that the perception or learning of statistical regularities is not performed by one neural region, but rather may be supported by multiple regions working in parallel (28, 32, 33, 39, 42, 43, for other hypotheses, see review 28). Sensory modality-general areas, such as the prefrontal cortex and the hippocampus, as well as lower perceptual or modality-specific regions, are proposed to subserve this capacity. However, detailed knowledge about the brain regions contributing to this dynamic and adaptive process is limited (3, 14, 29, 33, 39, 40, 44, 45).

To address this gap, we hypothesized that a core function of the brain is to encode TPs in a continuous and online fashion and that this is implemented in a distributed manner. Specifically, we investigated how different brain regions contribute to statistical learning by exploiting the high temporal and spatial resolution of intracranial electroencephalography (iEEG). We estimated the trial-by-trial information content of high-frequency activity (HFA; 75 to 145 Hz), a correlate of population neuronal spiking, from participants that were passively exposed to a sequence of randomly occurring tones. We then evaluated this information content estimate against the dynamic TPs of the sequence, stemming from an ideal observer model. Our results reveal that the brain continuously encodes the TPs in a stream of random stimuli through a network that spans areas outside the auditory system, including hippocampal, frontal, and temporal regions. Remarkably, this automatic process occurs even in the absence of evident relations within the stimuli or behavioral relevance.

## Results

### iEEG Unattended Listening Task

Participants (n=22; *Materials and Methods*) listened to a stream of tones where a standard tone alternated with deviant tones (P=0.5; inter-stimulus interval 500 ms). This stream followed a multi-dimensional auditory oddball paradigm, where deviant tones varied relative to the standard in terms of either frequency, intensity, perceived sound-source location, a shortened duration, or a gap in the middle of the tone (P=0.1 for each deviant type; Fig. 1, *Materials and Methods*). Within a set of ten tones (five standard tones and five deviant tones), each of the five deviant types was presented once in random order. For deviations in location, intensity, and frequency, two stimuli versions were used (P=0.5), namely, location left/right, intensity low/high, and frequency low/high. Together with the other two deviants, this resulted in eight potential deviants. During recording, participants were asked not to pay attention to the sounds while reading a book or magazine. All participants reported that they were able to focus on the reading material and did not attend to the tones or noticed any patterns in the stimuli.

**Figure 1:**
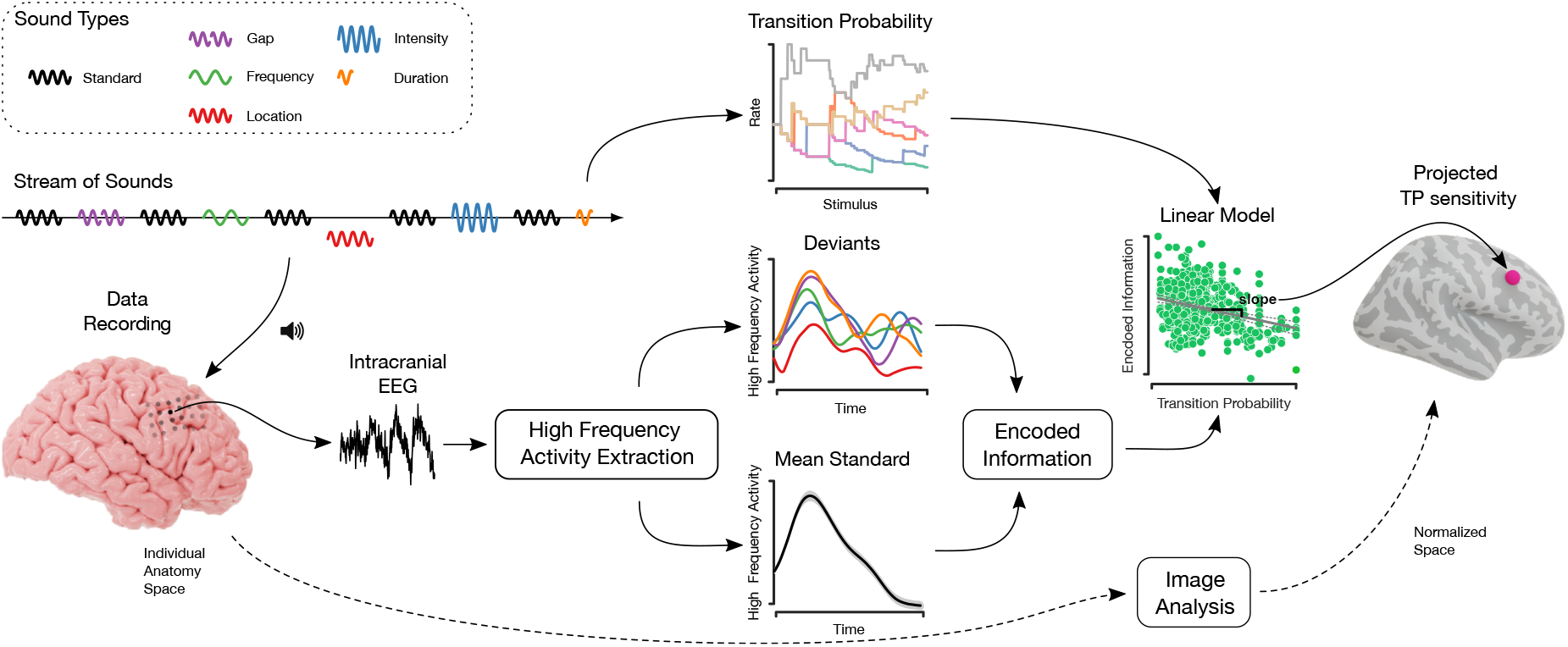
Overview of the analysis. An unattended listening task was presented to participants while recording their event-related electrical brain activity through intracranial electrodes. The emerging iEEG signal was then analyzed, resulting in HFA responses to standard and deviant tones. Based on the standards we computed a channel-specific mean standard response. Differences in normalized encoded information between deviant and mean standard responses were computed using a compression algorithm. The higher the value of this encoded information measure, the lower the similarity between the mean standard and a respective deviant tone response. In the next step, linear models between encoded information and TP estimates were employed, where the latter stemmed from an ideal observer analyzing the stream of sounds. After accounting for multiple comparisons, these channel-specific slopes were then projected onto the normalized anatomical space to enable comparison across subjects.

From the 22 participants, a total of 1078 channels (mean: 48, range: 12 to 104) were recorded. The recordings were manually cleaned by excluding noisy or epileptic channels or segments from the analysis. HFA was then reliably extracted from a total of 785 channels within cortical or subcortical structures, and HFA event responses (trials) were evaluated in the 400 ms time window following the sound onset.

### Encoded Information Peaks in Primary and Secondary Auditory Cortices

We estimated the information content of each deviant tone HFA response in relation to the HFA responses to standard tones. Accordingly, the information content in standard responses was used as a reference point to measure the information content in deviant responses, which yielded a normalized measure of *encoded information* for each deviant response (Fig. 1, bottom; *Materials and Methods*). Smaller values of *encoded information* suggest that the information content in deviants is similar to the one in standards, whereas greater values indicate a larger amount of encoded information in the responses to deviants compared to standards. To systematically evaluate the involvement level across the cortex, we defined regions of interest (ROIs) that typically engage in auditory processing and statistical learning tasks (1, 28, 32, 33, 41), comprising temporal, frontal, insular, peri-central sulci, and ACC cortices, as well as the hippocampus (Fig. 2a, Tab. S1). We then compared the *encoded information* across the ROIs. The greatest median *encoded information* values were observed in primary and secondary auditory cortices (superior temporal plane, insula posterior, and temporal lateral ROIs), suggesting that core aspects of deviant processing locate there (Fig. 2b, two-tailed pairwise Mann–Whitney–Wilcoxon tests, FDR corrected, p ≤1.12e-2, |z| ≥ 2.5). Each ROI’s median encoded information were significantly greater than zero (one-tailed Wilcoxon signed-rank test, FDR corrected, p ≤1.22e-4, z ≥ 4.53). Added together, these results indicate that the encoded information in the responses to deviants reflects the local sensitivity of specific brain areas to unexpected events in accordance with previous studies on deviance detection (46–49). Additionally, we examined the sensitivity to specific deviant types across ROIs. Statistical analysis only identified significant differences in the encoded information of specific deviant types in the superior frontal area. The statistically significant differences were between the deviant types of ”location left”, ”intensity up”, and ”frequency down” to ”gap”, respectively (Fig. S6, two-tailed pairwise Mann–Whitney–Wilcoxon tests, FDR corrected, p ≤5.30e-4, z ≥ 3.5).

**Figure 2:**
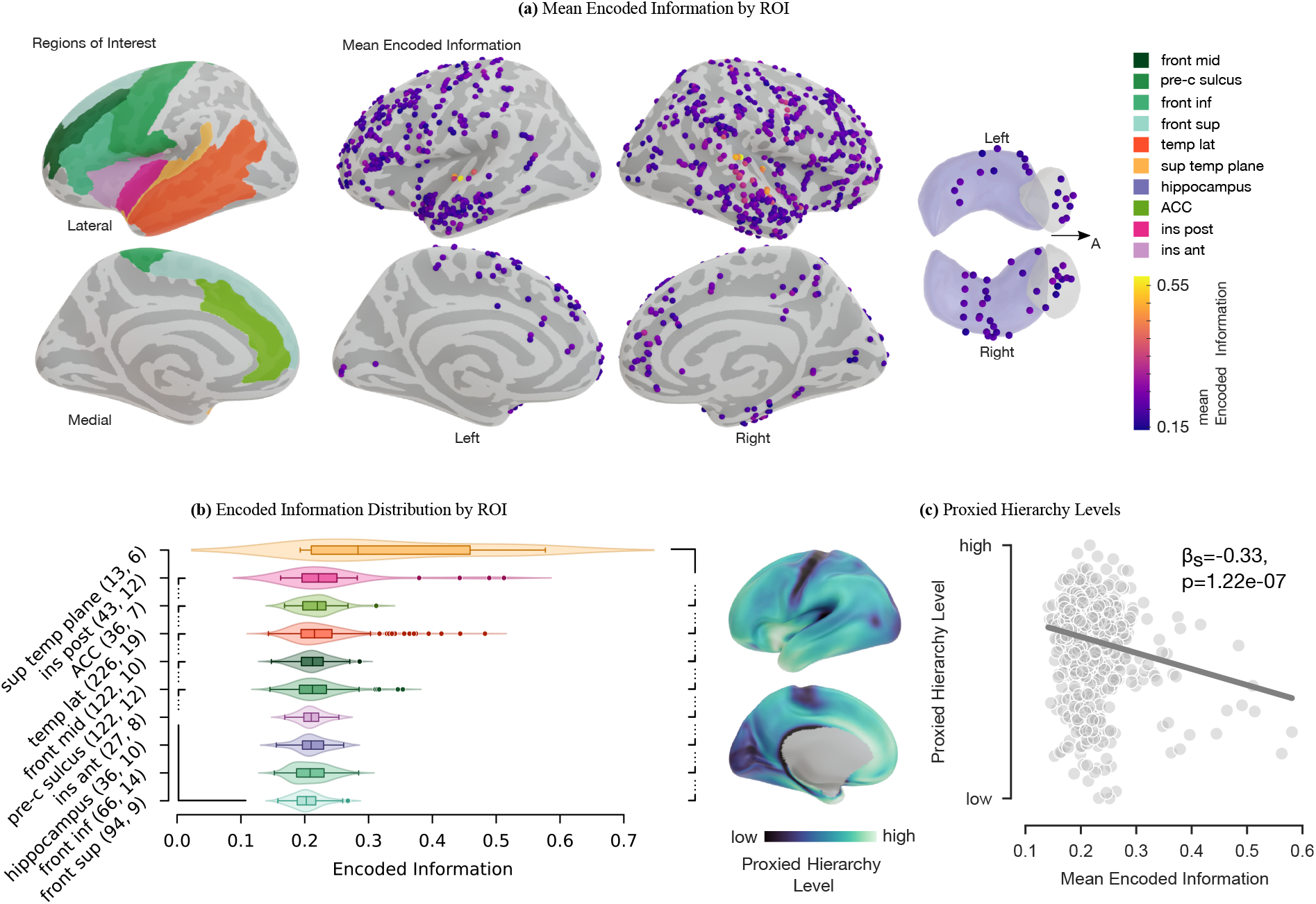
Illustration of the encoded information analysis results. **a**: Top: ROIs on the inflated brain model (Tab. S1 for full region labels). Bottom: lateral and medial view of the mean encoded information distribution across 22 subjects projected onto the inflated brain model. The image on the bottom right shows the transverse plane of the amygdala (gray) and hippocampus (purple). ”A” stands for the anterior direction. Each sphere represents one channel. **b**: Distribution of the ROIs’ encoded information. The number of channels (first) and subjects (second) for each ROI are in the axis labels. The nested brackets indicate a significant difference between median values.

### Encoded Information is Hierarchically Organized

Previous animal and human studies indicate a hierarchical organization of brain regions behind the detection of unexpected events (6, 41, 42, 50). We utilized a proxy measure of anatomical hierarchy to investigate to what extent this is reflected in the *encoded information* values across brain regions. Anatomical hierarchy can be defined as a global ordering of cortical areas corresponding to characteristic laminar patterns of inter-areal feedforward and feedback projections (5, 51, 52). Proxied cortical hierarchy levels that quantify these projections across the cortex were obtained from open-access structural magnetic resonance imaging (MRI) datasets from the S1200 subject release (53; *Materials and Methods*). Methodological constraints in (53) precluded the mapping of the hippocampus in the present analysis. Areas lower in the hierarchy (with predominantly feedforward projections) are primarily associated with primary sensory functions, whereas areas higher in the hierarchy are associated with higher cognitive functions (5, 51, 52). For each contact point, hierarchy level channel estimates were determined by taking the average value of all proximal points located within the contact point vicinity. We observed a significant negative correlation between the *encoded information* and the proxied cortical hierarchy levels (Fig. 2c; linear mixed-effects model with random effects for subjects: *y* = *β*_0_ + *β*_1_*x* + *b*_0_ + *ϵ* with proxied hierarchy level *y*, the encoded information *x*, the random effect for subjects 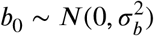 and the observation error 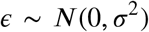; *β*_0_ = 0.16, 95% CI [0.13, 0.19], *β*_1_ = −0.33, 95% CI [−0.21 −0.44], p_*β*_1__=1.22e-7, *σ_b_*=3.94e-2, 95% CI [2.79e-2, 5.64e-2], *ϵ*=7.54e-2, 95% CI [7.16e-2, 7.92e-2]).

### Ensemble Activity Exhibits Sensitivity to Transitional Probabilities

During the time course of the stimuli, we incrementally estimated TPs in the fashion of an ideal observer. At each given deviant event (trial), TP estimates were updated based on all previously presented deviant stimuli. Consequently, TPs dynamically evolved along the course of the experiment since a finite stream, as opposed to an infinite horizon stream, naturally entails temporal patterns because of the alternating occurrence of deviants (Fig. 1, TP graph, *Materials and Methods*). To determine which brain area exhibits a sensitivity to these temporal relations we evaluated the relationship between HFA encoded information and the TPs of deviant tones through robust linear models. Before regression, the trial-specific encoded information values were normalized by the channel means to correct the encoded information that solely reflects auditory sound processing mechanisms (Fig. S7). For each channel, the resulting slope is defined as the channel-specific *TP sensitivity. TP sensitivity* values acted as an indicator of how sensitive the brain tissue around the channel was towards TPs in the stream of tones. Zero value *TP sensitivity* of a channel indicates that the encoded information in the deviant responses is not affected by the TPs of the events, whereas lower values imply a higher impact. Fig. 3a shows two example electrodes of high and low *TP sensitivity* (each green dot represents a deviant trial). In the analysis, 61.53 % of the 785 channels across all subjects showed a significant *TP sensitivity* (Fig. 3b & S3, permutation-based test, FDR corrected). These channels tended to increase the amount of *encoded information* in the HFA response when the likelihood of an event occurrence decreased (low TP) and conversely decreased the *encoded information* for more expectable events (high TP). Notably, the *TP sensitivity* distributes over the brain (Fig. 3c). Therefore we evaluated this distribution in terms of ROIs. Each ROI’s *TP sensitivity* except for the ACC were significantly lower than zero (one-tailed Wilcoxon signed-rank test, FDR corrected, p ≤2.97e-2, |z| ≥ 1.89), indicating that most ROIs were involved in the encoding of TPs (Fig. 3d). Importantly, our results were consistent across participants. Out of the 22 subjects, an average of 52.10 % (95% CI [47.19%, 57.04%]) showed a significant *TP sensitivity* across the ROIs (Fig. 3b & S4). Moreover, we studied differences in the *TP sensitivity* across ROIs, where hippocampus and inferior frontal cortex showed the greatest sensitivity to TPs (Fig. 3e, two-tailed Mann–Whitney–Wilcoxon tests, p ≤4.37e-2, |z| ≥ 2.02).

**Figure 3:**
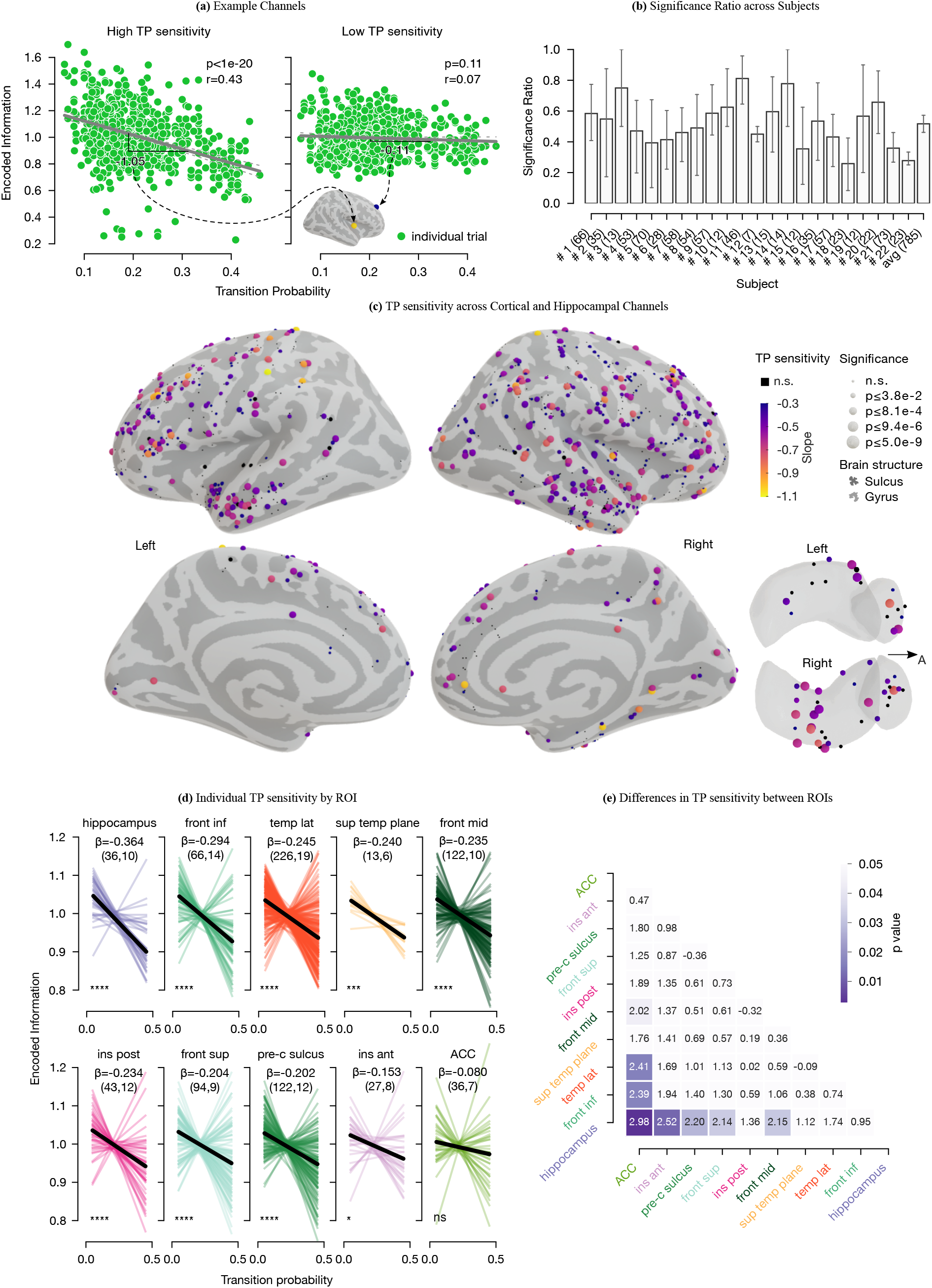
TP sensitivity results. **a**: Two example channels resulting from the robust linear regression between TPs and encoded information (each green dot represents a trial). For each channel, the encoded information values were normalized by their mean. The resulting slope indicates how sensitive a region underneath a contact point is towards the variation of TPs. The first channel shows a negative slope of −1.05. Thus, the more frequent a transition, the more the information encoded in the deviant response decreases. **b**: Ratio of the significant to total channels (number in brackets) across subjects. The error bars indicate the 95% CI across ROIs. **c**: Inflated brain model with lateral and medial views of the right and left hemispheres and a superior view of the amygdala and hippocampus. Each sphere represents a channel projected onto the surface. The colors indicate its TP sensitivity. TP sensitivities greater than −0.3 or within the first 25 % of all values have the lowest color in the color gradient. The size of the spheres indicates the p-values of the slopes. They are divided such that each interval contains 1/4 of the p-value set. **d**: TP sensitivity by ROIs, where the individual TP sensitivities (regardless of significance) are colored. In black, the median TP sensitivity is shown (see *β* for its numerical value). The number of channels and subjects are given in parentheses in the subtitles. Except for the ACC, all ROIs show a significant median TP sensitivity (statistical significance of the slopes is indicated with ”ns” p>0.05, * p≤5e-2, *** p≤1e-3, and **** p≤1e-4). **e**: Matrix of z-values representing individual statistical differences of TP sensitivity between ROIs.

## Discussion

We studied how humans passively listening to a multi-feature sequence of random sounds implicitly encode conditional relations between sounds. Crucially, our results show that the auditory system embedded in a distributed hierarchical network continuously monitors the environment for potential saliency, maintaining and updating a neural representation of temporal relationships between events. This suggests that the brain continuously attempts to predict and provide structure from events in the environment, even when they are not behaviorally relevant and have no evident relation between them.

Participants demonstrated remarkable sensitivity to TPs. From a statistical learning perspective (defined as “all phenomena related to perceiving and learning any forms of patterning in the environment that are either spatial or temporal in nature” (54)), these findings suggest an implicit learning process in which TPs are internally inferred. On average, more frequent deviant transitions exhibited less encoded information in the HFA responses. Conversely, rarer transitions showed an increase in the encoded information (Fig. 3a & 2c). Consequently, these results indicate an encoding of TPs, consistent with previous studies using more structured and stationary stimuli in humans and non-humans (4, 11–23, 25, 41, 42, 45). In our study, we additionally point out that the brain also is sensitive to dynamic TP courses in a randomly structured sequence of varied auditory stimuli. The brain’s sensitivity to TPs within our random sequence suggests a more general mechanism that continuously encodes TPs between events in the environment. This critical mechanism forms the basis of a statistical learning system wherein the brain integrates every event into an internal representation of the environment based on the statistical relationship between events. Since *a priori* the presence of patterns within stimuli is unknown, the brain might automatically encode their TP to detect potential structure and violations of such. Artificial grammar learning studies, where subjects learn patterns of nonsense words, confirm the relevance of this TP encoding in language learning (16, 28, 29, 33, 55).

Following the notion of predictive coding, the encoded information in each deviant response can be interpreted as a bottom-up prediction error signal, i.e., the amount of information in each novel event not explained away by top-down prediction signals (5, 7, 42, 56). Consequently, low TP events, i.e., less expected events, elicited a higher amount of encoded information and hence larger prediction errors derived from less accurate predictions. Accordingly, this information is used in higher cortical areas to update internal models for future predictions. On the other hand, high TP events, i.e., more expected events, elicited a lower amount of encoded information. This generates smaller prediction error signals and smaller updates of the internal models. Internal representations of TPs between events are fundamental to build useful predictions of upcoming events rather than simpler frequentist representations (12, 24). However, there is a lack of studies investigating TPs in predictive processing in general, while in statistical learning, there is a need for more neurophysiological studies. Our study takes a step forward in both of these directions, and shows that TPs might constitute a central statistic used by internal perceptual models at the core of predictive processing and statistical learning.

Our results provide novel evidence that the encoding of acoustic deviant transitions is anatomically distributed and not exclusively concentrated in auditory cortices (Fig. 3c). The automatic process of identifying temporal relationships is subserved by a network consisting of the hippocampus in concert with the inferior frontal, temporal, and insular cortices. Accordingly, by entailing multiple active brain regions, this network bundles findings from various prior statistical learning (28, 32) and predictive processing (6, 41) studies together.

Specifically, the hippocampus contributes most to temporal transition encoding between salient events. In contrast to other areas, hippocampal responses indicate high sensitivity to TPs while having a lower sensitivity to deviant tones (Fig. 2b & 3d). Accordingly, hippocampal activity may reflect a more generic context sensitivity to the events’ probabilistic structures, i.e., learning about event occurrences within a given structure itself instead of encoding actual deviating events (57). Our results provide new evidence for the role of the hippocampus during implicit learning, consistent with recent suggestions that this area is a rapid supramodal learner of arbitrary or higher-order associations in the sensory environment (3, 16, 28, 32, 33, 39–41, 45, 58, 59). In a recent iEEG study presenting 12 syllables within an auditory stream, Henin et al. (16) observed that TPs are encoded in lower-order areas of the superior temporal plane and not in the hippocampus, which uniquely represented the identity (i.e., the specific higher-order chunk such as a word) of their sequences. Therefore, the hippocampus did not appear to engage in forming the neural representation of TPs but performed operations that built upon them. We, on the contrary, found the hippocampus to be the main contributor among the cortical areas in encoding TPs. These differences might emerge because our study used passive listening with pure tones, while Henin et al. used active listening with syllables. Our results fit well with previous studies indicating the hippocampus’ fundamental role in statistical learning and encoding stimuli uncertainty, both attended and unattended (3, 16, 28, 32, 33, 39–41, 43, 45, 57–60). According to that, the hippocampus might operate differently depending on task demands. By its domain-general learning mechanisms, possible hippocampal involvement could comprise indirect modulation of lower-level sensory areas or direct computations of hippocampal representations (28, 32).

We also observed sensitivity to transitions between events in the inferior frontal cortex. Evidence of inferior frontal involvement in statistics-driven learning processes is sparse (28, 33, 41) and mainly relies on explicit learning studies using fMRI (8, 44). However, it is commonly described in the deviance detection literature, where a role of a higher hierarchical node is attributed to this region (46, 48, 49). Evidence from non-human primates iEEG studies manipulating the predictability of events also supports this involvement by showing a spatially dispersed contribution of regions that includes the prefrontal cortex in both passive auditory (42) and active visual paradigms (6).

Notably, channels in the superior temporal plane showed the highest encoded information and a high TP sensitivity (Fig. 3d), suggesting a key role of the supratemporal plane in both the deviance detection and the implicit learning of transitions between salient auditory events. This is consistent with previous reports about this region being active in conditional statistical learning (17, 33, 44, 45, 61). Thus, perceptual processing of individual stimuli in low hierarchical areas might be strongly affected by learning temporal patterns in streams of stimuli (22, 23, 28, 62). This is possibly due to a local process, top-down modulations, or both. However, previous studies have shown that top-down information flow interacts with bottom-up information flow at all levels of the hierarchy (5, 6, 48, 49).

An unexpected observation was the significant TP sensitivity of individual channels in the occipital lobe, indicating a contribution to TP encoding of the auditory stimuli. It has been shown that during auditory oddball and statistical learning paradigms, attentional processing can activate visual processing regions, which are typically engaged in the perception of visual objects (16, 63, 64). When queried, all of our participants reported that they could focus on the reading material and did not pay attention to the tones. Hence, this leaves open whether this auditory occipital activation might also be observable during passive listening tasks and whether this is specific to the sensitivity of our HFA recording. Current evidence is sparse, but two previous studies on deviance detection during passive listening showed similar occipital effects using fMRI and scalp EEG (64, 65).

In terms of deviance detection, our results suggest a main involvement of the superior temporal plane and posterior insula (Fig. 2b). Previous studies on auditory deviance detection using iEEG, MEG/EEG source localization, and fMRI have shown similar responses to deviants over the supratemporal plane (1, 34, 47–49, 65–70), but detailed information for the insular cortex is sparse. In line with recent reports about its contribution to auditory processing (66, 71), we found that the posterior part showed larger encoded information than the anterior part. We also noticed that the ACC, middle frontal and pre-central sulcus moderately engaged in change detection. Although not often observed in auditory experiments, activation of these regions has been previously reported in the context of pre-attentive oddball paradigms with frequency (or duration) deviants using EEG (65, 65, 68, 72) or fMRI (64, 70). In our study, the ACC contributes to auditory change detection but did not reach a significant sensitivity to TP, generally consistent with previous reports (65, 72). It is presumably more involved in cognitive control or error detection, such as recognizing global patterns (47, 67). In our pre-attentive paradigm, we speculate that the ACC monitors the high-level structure of individual deviant occurrences rather than the automatic TP encoding. Further, areas lower in the hierarchy are more sensitive to deviant tones, and conversely, higher hierarchy locations exhibit lower encoded information values (Fig. 2c). Interestingly, our results indicate that the encoding of deviants was not strictly confined to specific areas, but distributed across multiple brain regions in a hierarchically organized manner. This suggests that lower hierarchical levels, which show a preferential representation of the stimuli, are more sensitive to the different deviant tones. Together, these results are in line with studies on the hierarchical visual pathway which indicate that expectation suppression scales positively with image preference (73).

In our present study, we focused on the analysis of HFA, given that it captures fast fluctuations in iEEG. Aside from HFA, it might be especially worthwhile to consider lower frequency bands (e.g., alpha or beta) because these bands presumably carry information of predictions (5, 6). However, because iEEG represents the population activity of spiking neurons, concerning lower and thus less fluctuating frequencies, iEEG macroelectrodes may miss less prominent activity patterns of a minority of neurons (1).

Our work provides a comprehensive picture of neural correlates of statistical learning, which, before, were bundled together from multiple studies (28, 33, 45). Additionally, our setup shares similarities in common with language learning studies. Yet, the implications of our findings may be limited because our paradigm is implicit and employs pure tones. One possibility to account for this is to replace pure tones with syllables or chunks of sounds. Also, given the presumably different roles of brain regions during implicit and active learning tasks (16, 28), active exposure to our sound train could potentially allow a more direct comparison between brain regions, or to language learning studies.

Having ascertained implicit learning analytically through algorithmic information theory and having determined neural substrates that imply a cortical network of brain regions, we are now in the position to explore its underlying mechanisms and regional influences further. Specifically, adding lower frequency bands to our analysis would enable us to disentangle the distinct roles in information encoding and predictability signaling of sensory inputs. While having a lower HFA, evoked responses to predictable events might exhibit a higher alpha or beta activity (5, 6, 42). Accordingly, in the case of more frequently occurring, and thus more predictable transitions, there might be an alternative cascade of involved regions anchored in higher cortical areas. In that respect, it might be especially worthwhile to evaluate the preonset sound interval of event responses, phase-amplitude coupling or connectivity across ROIs.

Taken together, direct brain recordings reveal continuous encoding of structure in random stimuli. While automatically assessing the deviance of events, the brain simultaneously identifies patterns by encoding conditional relations between events, supporting both statistical learning and predictive coding frameworks. This implicit process involves, in addition to the hippocampus, inferior frontal cortices, pure sensory areas, and other cortical regions.

## Methods

### Stimuli

An unattended listening task following a multi-dimensional auditory oddball paradigm was used (48, 49, 74). The task consisted of a standard and five different deviant tones (Fig. 1). Standards had a duration of 75 ms with 7 ms up and down ramps and consisted of three sinusoidal partials of 500, 1000, and 1500 Hz. Deviants varied relative to the standard in the perceived sound-source location (left or right), intensity (±6dB), frequency (550, 1100, and 1650 Hz or 450, 900, and 1350 Hz), gap (25 ms silence in the middle), or by a shortened duration (^1^/3 or 25 ms shorter). Thus there were two stimuli versions for location, intensity, and frequency deviants. During the sequence, each standard tone was followed by a deviant tone. The deviant tone type was set up such that within a set of five consecutive deviants, each of the five types was presented once. In consecutive sets, the same deviant type did not repeat from the end of one set to the beginning of another. For the three deviants that had two stimuli versions, each version occurred equally often (P=0.5). Except for deviants varying in duration, all tones had a duration of 75 ms and were presented every 500 ms in blocks of 5 min consisting of 300 standards and 300 deviants. At the beginning of each block, 15 standards were played. To capture automatic, stimulus-driven processes, participants were asked not to pay attention to the sounds while reading a book or magazine. They completed 3 to 10 blocks, providing at least 1800 trials. Tones were presented through headphones using Psychtoolbox-3 (75).

### Participants

We recorded data from 22 (self-reported) normal-hearing adults with drug-resistant epilepsy who were potential candidates for resective surgery of epileptogenic tissue (mean age 31 years, range 19 to 50 years, 6 female). Patients underwent invasive intracranial electrocorticography (ECoG) or stereoelectroencephalography (SEEG) recordings as part of their pre-surgical evaluation. Intracranial electrodes were temporarily implanted to localize the epileptogenic zone and eloquent cortex. The number and placement of electrodes were guided exclusively by clinical requirements. Data were collected at El Cruce Hospital (n=15) and Oslo University Hospital (n=7).

### Data Acquisition

Pre-implantation structural MRI and post-implantation CT scans were acquired for each participant. ECoG or SEEG data were recorded using an Elite (Blackrock NeuroMed LLC, USA), a NicoletOne (Nicolet, Natus Neurology Inc., USA), or an ATLAS (Neuralynx, USA) system with sampling frequencies of 2000, 512, and 16 000 Hz, respectively.

### Electrode Localization

Post-implantation CT images were co-registered to pre-implantation MRI images using SPM12 (76). MRI images were processed using the FreeSurfer standard pipeline (77), and individual cortical parcellation images were obtained through the Destrieux atlas (78). Electrode coordinates were obtained with the iElectrodes Toolbox (79). Anatomical labels were automatically assigned to each contact based on the Destrieux atlas using the aforementioned toolboxes and confirmed by a neurologist/neurosurgeon. Coordinates were projected to the closest point on the pial surface (within 3 mm) and then coregistered to a normalized space using surface-based spherical coregistration (80).

### Signal-preprocessing

Monopolar intracranial EEG recordings were visually inspected and channels or epochs showing epileptiform activity or other abnormal signals were removed. Signals from electrodes located in lesional tissue or tissue that was later resected were also excluded. Bipolar channels were computed as the difference between signals recorded from pairs of neighboring electrodes in the same electrode array. In our study, we refer to these bipolar channels as ”channels” Data were low-pass filtered at 180 Hz, and line noise was removed using bandstop filters at 50, 100, and 150 Hz. Data were then segmented into 2000 ms epochs (750 ms before and 1250 ms after tone onset) and demeaned. We manually inspected and rejected epochs after bipolar re-referencing. To eliminate any residual artifact, we rejected trials with an amplitude larger than 5 SD from the mean for more than 25 consecutive ms, or with a power spectral density above 5 SD from the mean for more than 6 consecutive Hz. An average of 35 % of the trials were rejected, resulting in an average of 1592 trials analyzed per patient (range 728 to 3723). Data were resampled to 1000 Hz. Pre-processing and statistical analysis were performed in Matlab using the Fieldtrip Toolbox (81) and custom code. To obtain the HFA, preprocessed data were bandpass filtered into eight consecutive bands of 10 Hz bandwidth ranging from 75 to 145 Hz. The Hilbert transform was then applied to each filtered signal to obtain the complex-valued analytic time series, and the modulus of these signals computed to retain the analytic amplitude time series. Trials were baseline corrected (−100 to 0 ms) for each frequency band, and then the bands were averaged, producing a single time series per trial. Finally, for each channel, all trial time series were divided by the standard deviation pulled from all trials in the baseline period. For more information, see (66, Chap. 2).

### Encoded Information

We estimated the information content of HFA responses by employing the concept of Algorithmic Information Theory. This theory anchors in Algorithmic Complexity or Kolmogorov Complexity (K-complexity). The K-complexity is the ultimate compressed version or minimum description length of an object, i.e., its absolute information content (82). If the minimum description length is short (long), an object is characterized as ”simple” (”complex”). Because it is not possible to compute the theoretically ideal K-complexity, it is often heuristically estimated, obtaining an upper-bound approximation. Possible estimation approaches are conventional lossless data compression programs, e.g., gzip (82, 83).

Based on the K-complexity, various metrics were derived. One instance is the Normalized Information Distance or its estimation counterpart, the Normalized Compression Distance (NCD). The NCD allows to compare different pairs of objects with each other and suggests similarity based on their dominating features (or a mixture of sub-features) (82, 83). For a pair of strings (*x, y*), the NCD(*x, y*) is defined as

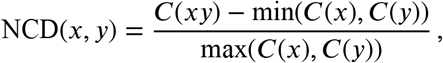

with *C*(*xy*) denoting the compressed size of the concatenation of *x* and *y*, and *C*(*x*) and *C*(*y*) their respective size after compression (82, 83). Further, the NCD is non-negative, that is, it is 0 ≤ NCD(*x, y*) ≤ 1+*ϵ*, where the *ϵ* accounts for the imperfection of the employed compression technique. Small NCD values suggest similar objects, and high values suggest rather different objects.

For each channel, we defined single-trial *encoded information* for each deviant response by computing the NCD measure between the HFA deviant response and the channel-specific mean HFA standard response (Fig. 1). Before their compression, HFA responses were represented by grouping their values into 128 discrete steps (bins). The bins covered equal distances and in a range between the global extrema of all trials considered. The compressor then received the indices of the bins that contained the elements of the signals (84, 85). Compression proceeded through a compression routine based on Python’s standard library and gzip. To account for the differences in auditory sound processing across channels the trial-specific encoded information values were normalized in terms of the channel mean of encoded information for the TP sensitivity analysis.

### Transitional Probability

We estimated conditional statistics describing the inter-sound relationship through TPs between adjacent deviant tones. After each deviant tone presentation (Fig. 1), TPs were determined through estimating their maximum-likelihood (14, 25, 26, 86), i.e., through

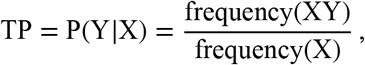

for each event-to-event combination X or Y. For each time step, resulting TPs were then stored in a TP matrix (stochastic matrix of size 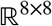.

### Anatomical Hierarchy

Human T1w/T2w maps were obtained from the Human Connectome Project (53). The maps were then converted from the surface-based CIFTI file format to the MNI-152 inflated cortical surface template with Workbench Command (87). The structural neuroimaging maps are suggested to be a measure sensitive to regional variation in cortical gray-matter myelin content (51). One function of myelin might be to act as an inhibitor of intra-cortical circuit plasticity. Early sensory areas may require less plasticity, hence more myelination, and hierarchically higher association areas, in turn, have less myelination, presumably enabling greater plasticity (88). Accordingly, T1w/T2w maps may serve as a non-invasive proxy of anatomical hierarchy across the human cortex through an inverse relationship. The anatomical hierarchy can be defined as a global ordering of cortical areas corresponding to characteristic laminar patterns of inter-areal projections (5, 51, 52). To directly work with the hierarchy ordering, T1w/T2w maps were inverted and normalized to the value range of our data set.

### Statistical Analysis

For the statistical analysis, the first 30 trials of each recording block were disregarded. By that we aimed to exclude the initial phase of the experiment that potentially biases our correlation analysis. To estimate the TP sensitivity of a channel, the eight distinct tone types were grouped into one regressor. Subsequently, robust linear regression was performed in Matlab (Fig. 1 & 3a), where TP values greater than 0.7 were excluded. For the regression, an alpha value of 0.05 was considered significant. To correct for multiple comparisons, false discovery rate (FDR) adjustment was applied with an FDR of 0.05. Further, our linear regression model examined the relationship between information content and TPs of all adjacent deviant transitions. For this reason, we performed surrogate data testing for uncorrelated noise on the regression models by building shuffled surrogates of the regressor variables encoded information and TP (Fig. S3).

## Ethics Approval and Consent to Participate

This study was approved by the Research Ethics Committee of El Cruce Hospital, Argentina, and the Regional Committees for Medical and Health Research Ethics, Region North Norway. Patients gave written informed consent prior to participation.

## Data Availability

The study in this article earned Open Materials for transparent practices. Materials for the experimental scripts and stimuli, and custom analysis code is available at. Materials for the experimental scripts and stimuli, and custom analysis code is available at osf.io/2n6c9. Due to the confidential nature of the data, the patients’ datasets analyzed for the current study are not publicly available. Our ethical approval conditions that do not permit public archiving of study data. Readers seeking access to the data supporting the claims in this paper should contact the corresponding author Alejandro Blenkmann, Department of Psychology, University of Oslo; the Research Ethics Committee of El Cruce Hospital, Argentina; and the Regional Committees for Medical and Health Research Ethics, Region North Norway. Requests must meet the following specific conditions to obtain the data: a collaboration agreement, data sharing agreement, and a formal ethical approval.

## Acknowledgments

We thank the patients for kindly participating in our study. We want to express our gratitude to the EEG technicians at El Cruce Hospital and Oslo University Hospital-Rikshospitalet for their support. We thank Yamil Vidal, Fernando Rosas, and RITMO colleagues for rich discussions. This work was partly supported by the Research Council of Norway (RCN) through its Centres of Excellence scheme project number 262762, RCN project number 240389 and 314925, NINDS Grant R37NS21135, NIMH CONTE Center P50MH109429, and Brain Initiative U01-NS108916.

## Author contributions

JF, AOB, AKS, and TE designed this study. AOB carried out the experiment and collected the data. JF and AOB performed the analyses. TE, AKS, SK, and TB provided the data. JI implanted and reviewed depth electrodes. JF, AOB, KG, TE, AKS, and RTK contributed to the interpretation of the results. JF wrote the manuscript with inputs from AOB, KG, TE, and RTK. All authors revised the manuscript. All authors read and approved the final manuscript.

