## Supplementary Information for "Direct brain recordings reveal continuous encoding of structure in random stimuli"

In this document, first, a detailed description of the regions of interests (ROIs) is supplied. Second, the significance testing by use of surrogates is demonstrated (Fig. S2). Then, statistically significant differences of the encoded information measure across ROIs is presented (Fig. S1), followed by the significance ratio (number of significant to total channels) across ROIs and subjects (Fig. S3). In Fig. S4, the distribution of the Pearson correlation coefficients is demonstrated. Subsequently, the deviant-specific encoded information across ROIs is depicted (Fig. S5), and, lastly, the effect of the normalization step is visualized (Fig. S6).

#### Contents

**1 Regions of Interest** 2

**2 Mean Encoded Information across ROIs** 3

**3 Significance Testing** 4

**4 Significance Ratio Across ROIs** 5

**5 Pearson Correlation Coefficient** 6

**6 Deviant-specific Encoded Information across ROIs** 7

**7 Normalization of encoded information for TP sensitivity** 8

**References** 8

### 1 Regions of Interest

**Table S1:** Anatomical parcellations of the ROIs after (1).

| ROI | Short name | Long name |
|---|---|---|
| Superior temporal plane | G_temp_sup-Lateral | Lateral aspect of the superior temporal gyrus |
| | G_temp_sup-G_T_transv | Anterior transverse temporal gyrus (of Heschl) |
| | G_temp_sup-Plan_tempo | Planum temporale or temporal plane of the superior temporal gyrus |
| | S_temporal_transverse | Transverse temporal sulcus |
| Lateral temporal cortices | G_temp_sup-Lateral | Lateral aspect of the superior temporal gyrus |
| | G_temporal_inf | Inferior temporal gyrus |
| | S_temporal_sup | Superior temporal sulcus (parallel sulcus) |
| | S_temporal_inf | Inferior temporal sulcus |
| | G_temporal_middle | Middle temporal gyrus |
| Superior frontal cortices | S_front_sup | Superior frontal sulcus |
| | G_front_sup | Superior frontal gyrus |
| Middle frontal cortices | S_front_middle | Middle frontal sulcus |
| | G_front_middle | Middle frontal gyrus |
| Inferior frontal cortices | S_front_inf | Inferior frontal sulcus |
| | G_front_inf-Opercular | Opercular part of the inferior frontal gyrus |
| | G_front_inf-Orbital | Orbital part of the inferior frontal gyrus |
| | G_front_inf-Triangul | Triangular part of the inferior frontal gyrus |
| | Lat_Fis-ant-Horizont | Horizontal ramus of the anterior segment of the lateral sulcus (or fissure) |
| | Lat_Fis-ant-Vertical | Vertical ramus of the anterior segment of the lateral sulcus (or fissure) |
| Pre-central sulci | G_precentral | Precentral gyrus |
| | S_central | Central sulcus (Rolando's fissure) |
| | S_precentral-sup-part | Superior part of the precentral sulcus |
| | S_precentral-inf-part | Inferior part of the precentral sulcus |
| | G_and_S_paracentral | Paracentral lobule and sulcus |
| | G_and_S_subcentral | Subcentral gyrus (central operculum) and sulci |
| Anterior cingulate cortices | G_and_S_cingul-Ant | Anterior part of the cingulate gyrus and sulcus (ACC) |
| | G_and_S_cingul-Mid-Ant | Middle-anterior part of the cingulate gyrus and sulcus (aMCC) |
| Anterior insula | S_circular_insula_ant | Anterior segment of the circular sulcus of the insula |
| | S_circular_insula_sup | Superior segment of the circular sulcus of the insula |
| | G_insular_short | Short insular gyri |
| Posterior insula | S_circular_insula_inf | Inferior segment of the circular sulcus of the insula |
| | G_Ins_lg_and_S_cent_ins | Long insular gyrus and central sulcus of the insula |

#### 2 Mean Encoded Information across ROIs

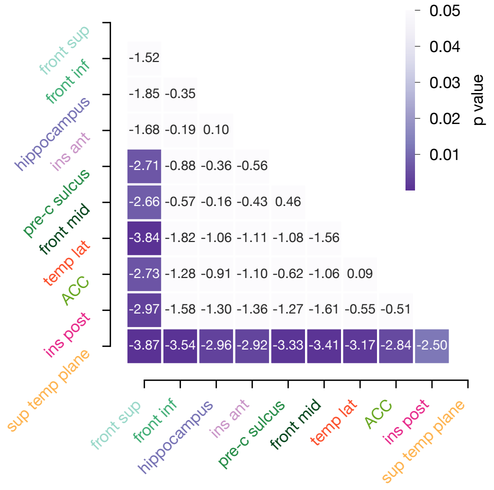


**Figure S1:** Matrix of z-values representing individual statistical differences of the encoded information measure (two-tailed pairwise Mann–Whitney–Wilcoxon tests) across ROIs.

### 3 Significance Testing

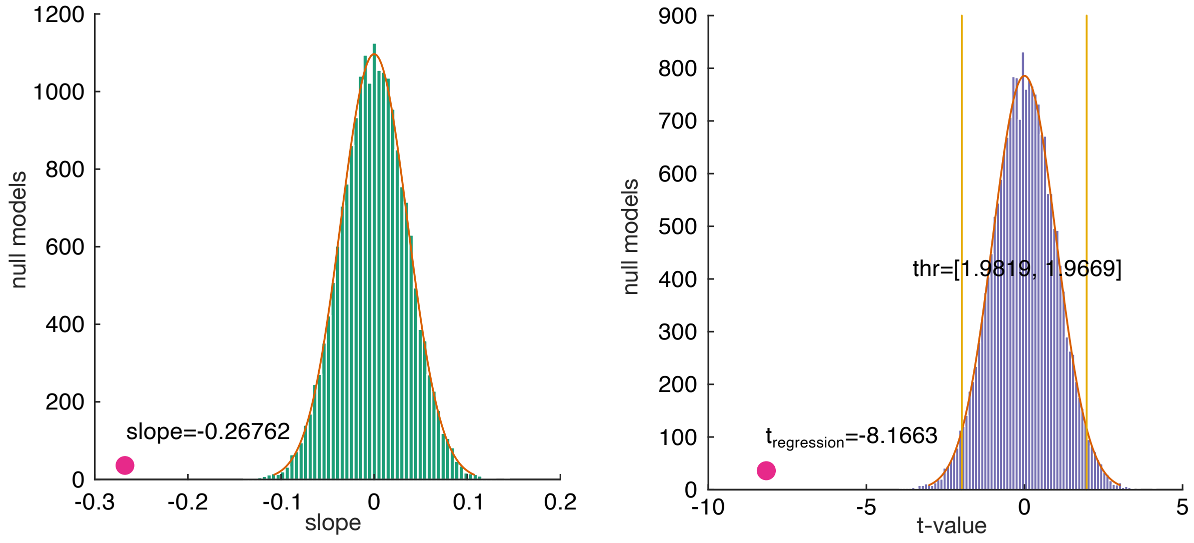


**Figure S2:** Significance testing through surrogates for an example channel. By randomly shifting the elements of the regression array that consists of encoded information and transition probalbities (TPs), 2*e*4 null models are generated. The hypothesis that the regression correlates noise can be rejected, when the respective t value is above or below the 2.5 % or 97.5 % quantile of the null model distribution (2).

### 4 Significance Ratio Across ROIs

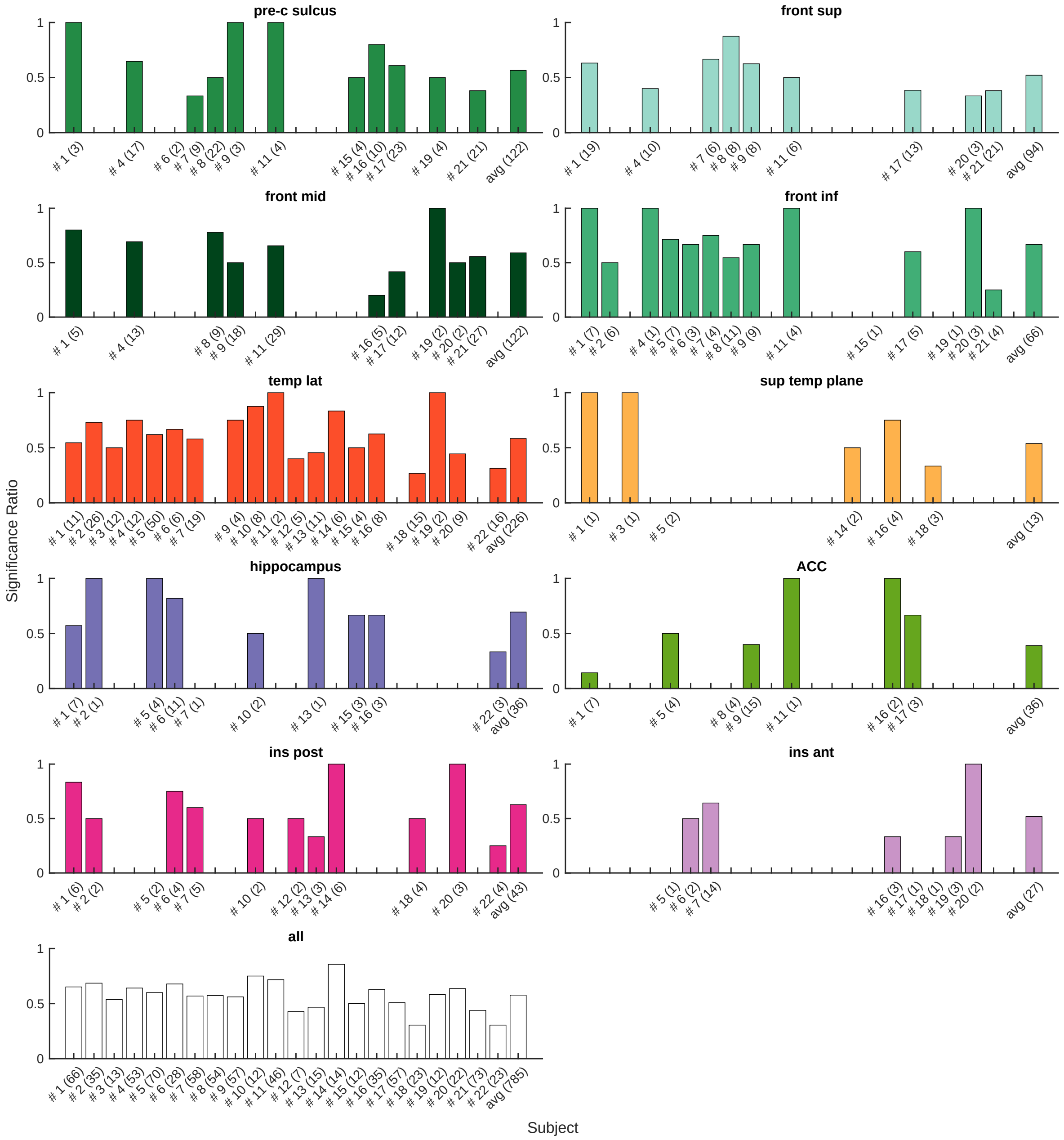


**Figure S3:** Distribution of the significant channels across subjects. 485 out of 785 channels show a significant slope after adjusting for channels-wise multiple correction and surrogate testing. On the x-label, the number in brackets shows the amount of channels each subject has in the respective ROI. The ratio itself is defined as the number of significant channels to total number of channels. On the bottom left, all ROIs are taken together.

### 5 Pearson Correlation Coefficient

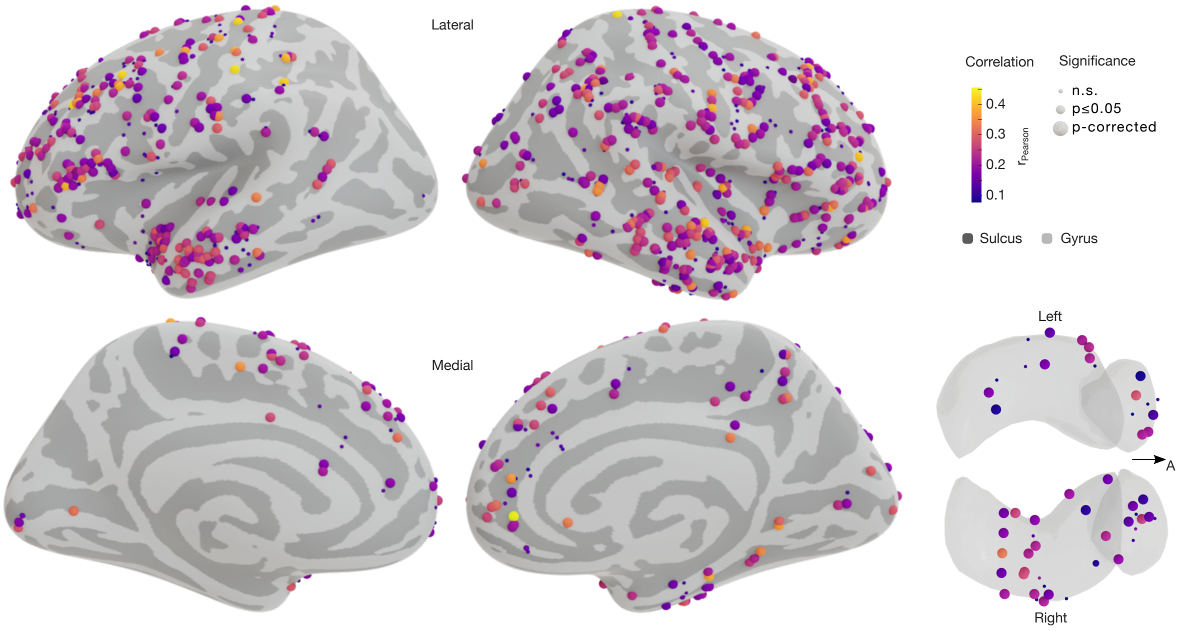


**Figure S4:** Inflated brain model with lateral and medial views of the right and left hemispheres and a superior view of the amygdala and hippocampus. Each sphere represents a channel projected onto the surface with the colors indicating its Pearson correlation coefficient resutling from the regressio of encoded information to TPs. The size of the spheres indicates the p-value corresponding to the performed regression. The p-values are divided such that each interval contains 1⁄4 of the p-value set.

### 6 Deviant-specific Encoded Information across ROIs

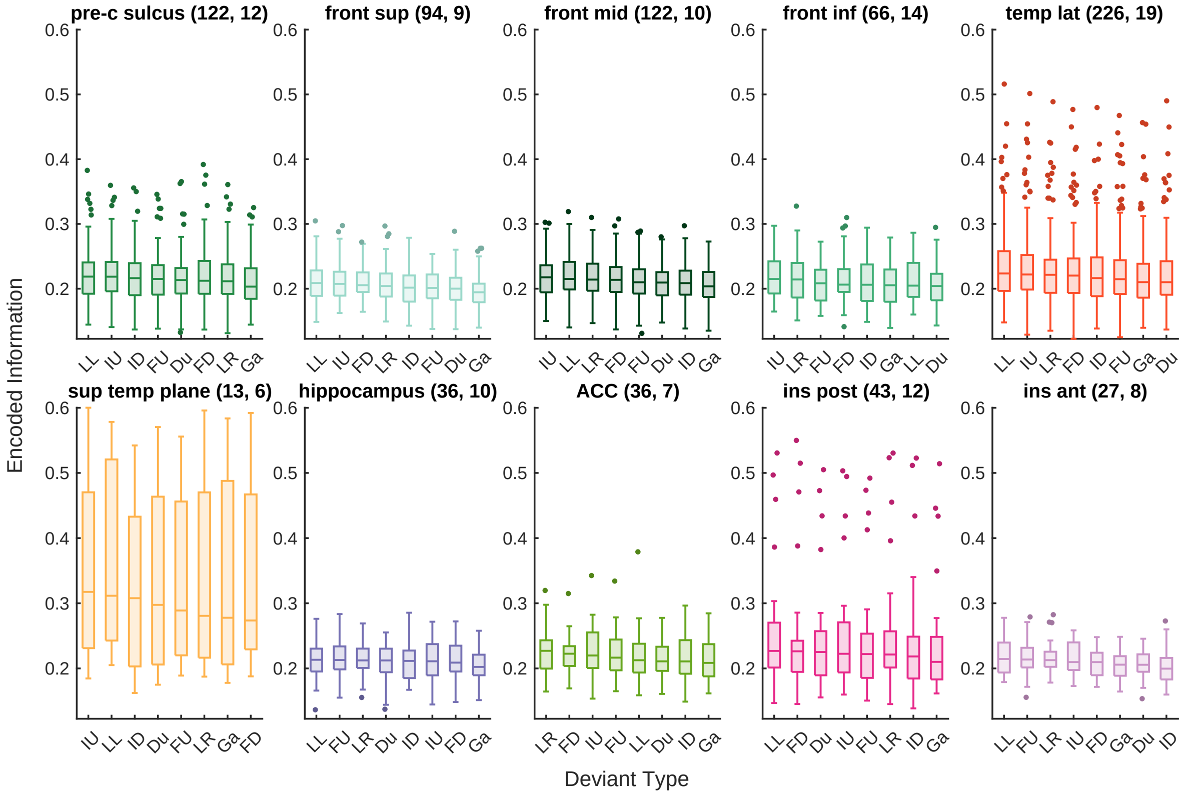


**Figure S5:** Distribution of the encoded information measure across ROIs for individual deviant types. In the axes labels ”LL/R” stands for location left/right, ”IU/D” for intensity up/down, ”FD/U” for frequency down/up, ”Du” for duration and ”Ga” for gap. In the titles, the term after each ROI name indicates the number of channels (first) and subjects (second). Statistical analysis showed no significant differences in the *encoded information* of specific deviant types in any of the areas but for superior frontal area with differences between the deviant types of ”location left”, ”intensity up”, and ”frequency down” to ”gap”, respectively (two-tailed pairwise Mann–Whitney–Wilcoxon tests, FDR corrected, p $\leq 5.30e{-4}$, z $\geq$ 3.5).

### 7 Normalization of encoded information for TP sensitivity

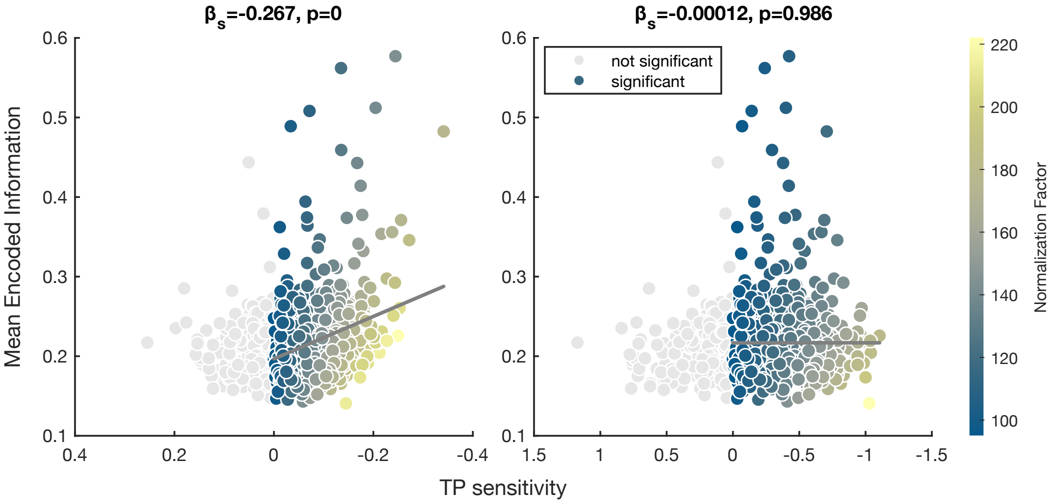


**Figure S6:** Relation between the mean encoded information and TP sensitivity before (left) and after (right) normalization. Without the channel-weise normalization of the encoded information measure, there is a significant positive correlation between the mean encoded information and TP sensitivity (left graph; linear mixed-effects model with random effects for subjects: $y = \beta_0+\beta_1 x+b_0+\epsilon$, with the mean encoded information $y$, the TP sensitivity $x$, the random effect for subjects $b_0 \sim N(0, \sigma_b^2)$ and the observation error $\epsilon \sim N(0, \sigma^2)$; $\beta_0 = 0.19$, 95% CI [0.19, 0.20], $\beta_1 = 0.28$, 95% CI [0.22 0.33], $p_{\beta_1} = 0$, $\sigma_b = 1.39\text{e} - 2$, 95% CI [9.46$e$–3, 2.04$e$–2], $\epsilon = 4.09\text{e} - 2$, 95% CI [3.89$e$–2, 4.29$e$–2]). As we are interested in the effect that solely can be traced back to the variance of the TPs, we adjusted for that by normalizing each encoded information value for each trial by its respective channel mean. Applying this step corrects for any correlation between mean encoded information and TP sensitivity (right graph). Consequently, by regressing encoded information with TPs, a signifiant slope only emerges if there is a real effect between the two regressors.
